# Permethylation of ribonucleosides provides enhanced mass spectrometry quantification of post-transcriptional modifications

**DOI:** 10.1101/2022.01.26.477959

**Authors:** Yixuan Xie, Kevin A. Janssen, Alessandro Scacchetti, Roberto Bonasio, Benjamin A Garcia

## Abstract

Chemical modifications of RNA are associated with fundamental biological processes such as RNA splicing, export, translation, degradation, as well as human disease states such as cancer. However, the analysis of ribonucleoside modifications is impeded due to the hydrophilicity of the ribonucleoside molecules. In this research, we used solid-phase permethylation to derivatize the ribonucleosides, and the permethylated ribonucleosides, which were then quantitively analyzed using a liquid chromatography−tandem mass spectrometry (LC−MS/MS)-based method. The solid-phase permethylation efficiently derivatized the ribonucleosides, and more than 60 RNA modifications were simultaneously monitored using ultrahigh-performance liquid chromatography coupled with triple quadrupole mass spectrometry (UHPLC-QqQ-MS) performed in the dynamic multiple reaction monitoring (dMRM) mode. Because of the increased hydrophobicity of permethylated ribonucleosides, this method enhanced retention, separation, and ionization efficiency, resulting in improved detection and quantification when compared to existing analytical strategies of RNA modifications. We applied this new approach to measure the extent of cytosine methylation and hydroxymethylation in RNA obtained from mouse embryonic stem cells with genetic deficiencies in ten-eleven translocation (TET) enzymes. The results matched previously performed analyses and highlighted the sensitivity, efficacy, and robustness of the new method. The advantage of this method enables comprehensive analysis of RNA modifications in biological samples.

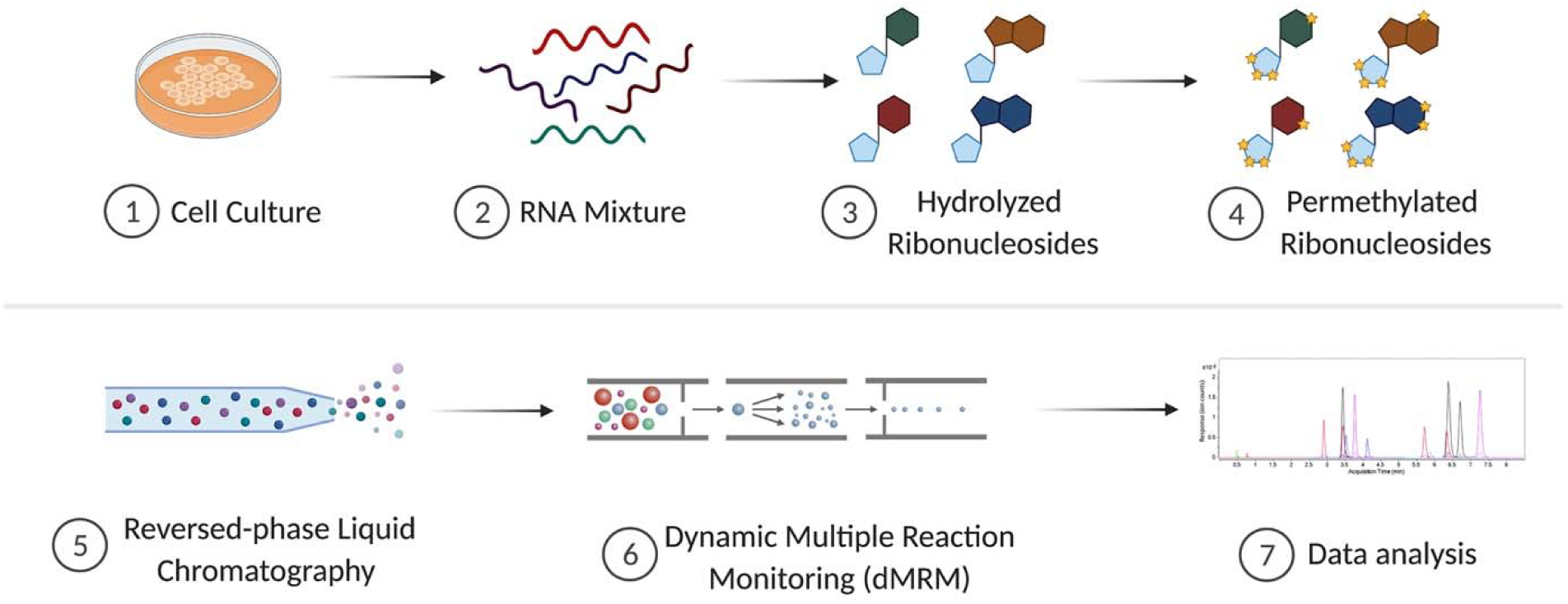

## Introduction

Ribonucleic acids (RNAs) are at the heart of the central dogma of biology. Messenger RNAs (mRNAs) are transcribed from the genome and direct protein synthesis in the cytosol. This process is highly regulated and aided by a number of noncoding RNAs, including transfer RNAs (tRNAs) and ribosomal RNAs (rRNAs). In recent years, it has become evident that chemical modifications deposited on RNAs during or after transcription contribute to the regulation of gene expression, giving rise to a field sometimes referred to as “epitranscriptomics”.^1-3^ Collectively, more than 150 distinct RNA modifications originating from the canonical adenosine (A), guanosine (G), cytidine (C), and uridine (U) have been identified across different organisms.^4^ These RNA modifications are involved in many fundamental biological processes, including cell differentiation, alternative splicing, and stress responses.^5^ At the molecular level, the modifications have been demonstrated to be crucial for proper RNA folding, topology, stability, high-order structure, and protein translation.^6, 7^ Translation is highly dependent on RNA modifications, which are critical for proper rRNA and tRNA function, and can regulate the rate and fidelity of translation when decorating mRNA.^8^ Additionally, RNA modification can affect the ability of proteins to recognize certain RNA.^9, 10^ Dysregulation of RNA modifications is associated with several diseases. For example, one of the most prevalent RNA methylations, N6-methyladenosine (m6A), has been linked to the carcinogenesis and progression of colorectal cancer.^11^

Although the importance of RNA modifications has been well-recognized, their characterization and quantification has been a long-standing problem hampered by the inadequacy of existing analytical methods. Sequencing-based techniques are powerful; however, most of the existing methods are unable to identify the modifications directly.^12^ While nanopore sequencing has been developed to directly analyze m6A, while this approach lacks sensitivity and it is challenging to detect other types of modifications.^13^ Indirect identification has been robustly performed using modification-specific antibodies and chemical reagents that react with specific modifications, but these methods cannot be used to make comprehensive and quantitative measurements because each modification requires a distinct workflow.^14^ Moreover, sequencing cannot reliably provide simultaneous quantitation and localization of multiple modifications. In addition to these genomic approaches, nuclear magnetic resonance (NMR) has been applied to RNA modification analysis.^15^ However, the analysis of modifications using NMR is challenging due to chemical shift overlaps, and line broadening often leads to high background noise and signal loss.^16^

Mass spectrometry (MS) has emerged as an alternative for developing powerful methods to analyze the modifications of RNA.^17^ Different modifications can be characterized in a single run, which enables high-throughput analyses. Although methods have been developed to analyze RNA at the oligonucleotide level to localize modifications, MS analysis of oligonucleotides is still far from established due to three limitations: digestion, instrumentation, and software.^18-20^ Therefore, hydrolyzing RNAs into ribonucleosides and characterizing RNA modifications at the ribonucleoside level using MS is a more accessible approach.^21, 22^ However, the commonly used reversed-phase chromatography is not optimal for native ribonucleosides due to their high polarity of most of these compounds. Although other stationary phases such as porous graphitic carbon (PGC) and hydrophilic interaction liquid chromatography (HILIC) have been utilized, these methods are not ideal for profiling all known ribonucleoside modifications,^23^ because these materials are not ideal for separating all modified ribonucleoside; for example, N6-threonylcarbamoyladenosine (t6A) cannot be reliably analyzed using PGC as a stationary phase.^24^ Additionally, some of these chromatography approaches for RNA analysis often use ion-pairing agents, which are not compatible with mass spectrometry and are difficult to remove from the liquid chromatography (LC).^25^

Rather than changing the material in the stationary phase, chemical derivatization methods have been applied to alter the chemical and physical properties of ribonucleosides and make them compatible with a more desirable stationary phase. Patteson *et al*. developed a method using 1-cyclohexyl-3-(2-morpholinoethyl)carbodiimide to label pseudouridine (Ψ) residues.^26^ Huang *et al*. also demonstrated a method to derivatize 5-methylcytosine (m5C) and its oxidation products using bromoacetonyl-containing reagents.^27^ These approaches were applied to exclusively monitor specific targets of interest depending on the selective chemical reactivities towards a particular ribonucleoside modification. Cai and co-workers converted ribonucleosides into acetonides using acetone for a more comprehensive study, and more than 50 derivatized ribonucleosides were identified.^28^ Nevertheless, the chemistry requires *cis–diol* groups, and nucleosides containing 2’-*O*-methylation could not be identified by this approach.

Here, we report a novel method to detect RNA modifications by MS that takes advantage of permethylation as a derivatization strategy. Permethylation replaces the hydrogens on hydroxyl groups, amine groups, and carboxyl groups with methyl groups. It is an efficient strategy for derivatizing a wide array of polar molecules and has been widely employed for glycan analysis.^29^ Permethylation of ribonucleosides were first attempted when the reaction was expected to yield volatile derivatives for gas chromatography–MS analysis.^30, 31^ As these studies using permethylation of canonical ribonucleosides were carried out before modern MS-based analytical methods were developed, the properties of the products were not well characterized using modern LC-MS. In addition, the unmodified ribonucleosides and the differently methylated ribonucleosides, such as A, m6A, and Am, cannot be unambiguously identified because the previous method only obtained MS1 spectra and lacked chromatographic separation. We therefore established a solid-phase permethylation method using isotopically labeled iodomethane to derivatize mono-ribonucleosides. The generated hydrophobic ribonucleoside derivatives were subsequently separated by reversed-phase C_18_-based liquid chromatography (RP-LC). Precursor and product ions were detected and quantified using triple quadrupole MS (QqQ MS) operated in the targeted dynamic multiple reaction monitoring (dMRM) mode, simultaneously. This method yielded a few advantages: (i) the solid-phase permethylation allowed efficient labeling in a short time period; (ii) the endogenous methylated ribonucleosides were distinguishable by different precursor and/or product ions due to derivatization with isotopically labeled iodomethane; (iii) the permethylation product of endogenous methylated and unmodified ribonucleosides had the same retention time, allowing improved the quantitative analysis; (iv) this method allowed spontaneous distinguishment for some ribonucleoside isomers (e.g., m3U and m5U) due to their unique precursor and product ions after permethylation; (v) the method yielded higher sensitivity (sub-femtomole level) compared to analyzing the underivatized ribonucleosides; and (vi) the chemical reaction was highly predictable, so that this method could be easily extended to currently unknown RNA modifications as they continue to be discovered. Consequently, more than 60 transitions were built as an example using the combination of ribonucleoside standards and digested ribonucleosides from cell extracts. To demonstrate the method’s viability, we examined 5-methylcytosine (m5C) levels in wild-type (WT) mouse embryonic stem cells (mESCs) and compared them to those found in cells deficient in enzymes of the ten-eleven translocation (TET) family. As previously reported, we observed the abundance of 5-hydroxymethylcytosine (hm5C) decreased in cells lacking one or more TET enzymes.^32^ In addition, our improved quantification method revealed an expected but previously unreported increase in m5C levels, whereas the level of 2’-O-methylcytosine (Cm) were similar in WT and mutant cells. This work demonstrates the advantage of permethylation as a derivatization strategy for highly accurate quantification of RNA modifications via mass spectrometry.

## Experimental procedures

### Samples and Materials

The ribonucleoside standards were purchased from Carbosynth (San Diego, CA). Iodomethane-d_3_, dichloromethane (DCM), zinc chloride (ZnCl_2_), sodium acetate (NaOAc), sodium hydroxide beads (NaOH), nucleosides test mix, nuclease P1, phosphodiesterase I, and phosphodiesterase II were purchased from Sigma-Aldrich (St. Louis, MO). Anhydrous dimethyl sulfoxide (DMSO) was purchased from Biotium (Fremont, CA). Acetonitrile (ACN), formic acid, recombinant shrimp alkaline phosphatase, porous graphitic carbon (PGC) stage tip, and micro spin column were purchased from Thermo Scientific (Waltham, MA).

### Culturing Mouse Embryonic Stem Cells (mESCs)

*Tet2* KO and *Tet1/2/3* triple KO mESC cell lines were previously described.^32^ All the cells were cultured on gelatin-coated dishes in KnockOut DMEM supplemented with 15% FBS, 0.1 mM MEM nonessential amino acids, 0.11⍰mM 2-mercaptoethanol, 1 mM L-glutamine, 0.5% penicillin streptomycin, 100⍰U/mL leukemia inhibitory factor (LIF), 31⍰μM CHIR99021, and 11⍰μM PD0325901, and the cells were maintained in a humidified cell culture incubator with 5% CO_2_ at 37 °C.

### Preparation of ribonucleosides from cell culture

Total RNA was extracted with Direct-zol RNA Kits (Thermo Scientific, Waltham, MA) based on the manufacture protocols. 100 ng of RNA samples were digested into ribonucleosides with 5 mU/μL of nuclease P1, 5 mU/μL of recombinant shrimp alkaline phosphatase, 500 μU/μL of phosphodiesterase I, and 6.25 μU/μL of phosphodiesterase II in 20 μL of digestion buffer (1 mM ZnCl_2_, 30 mM NaOAc, pH 7.5) overnight at room temperature. The digested ribonucleosides were purified using PGC stage tips and dried in a Savant SpeedVac concentrator (Thermo Scientific, Waltham, MA).

### Solid-phase permethylation of ribonucleosides

The digested ribonucleoside samples were permethylated using solid-phase permethylation, as previously described, with some optimization.^33^ Briefly, NaOH beads were packed into the empty spin columns (about 2 cm height), and the beads were washed with 100 μL of DMSO twice. The purified and dried ribonucleoside samples were reconstituted in a mixture of 1 μL of water, 50 μL of DMSO, and 30 μL of iodomethane-d_3_. The samples were loaded into the spin column and spun down at 200 x g, followed by reloading the samples into the column four times. Next, 20 μL of iodomethane-*d*_3_ was added to the sample and incubated at room temperature for another 10 min. The column was washed with 50 μL of DMSO twice, 500 μL of ice-cold water was added into the sample, and the mixture was incubated at room temperature for at least 1 min to quench the permethylation. Afterward, 300 μL of DCM was added, the liquid−liquid extraction was repeated at least five times for each sample, and the organic layer was dried using a Savant SpeedVac concentrator.

### Ultrahigh-Pressure Liquid Chromatography/Triple Quadrupole Mass Spectrometry (UHPLC/QqQ-MS) Analysis

Separation and characterization of the ribonucleosides were carried out on a Thermo Scientific Vanquish Flex binary UHPLC system coupled to a Thermo Scientific TSQ Altis QqQ mass spectrometer (Thermo Scientific, Waltham, MA). For the analysis, 2 μL of the sample was injected onto a Thermo Scientific Accucore Vanquish C18 column (150 × 2.1mm, 1.5μm) and separated using a 20 min binary gradient with a constant flow rate of 0.2 mL/min at 60 °C. Mobile phase A was water with 0.1% formic acid, and mobile phase B was 80% ACN/water (v/v) with 0.1% formic acid. To analyze the underivatized ribonucleosides, the following binary gradient was used: 0−3 min, 0% B; 3−10 min, 0-2% B; 10−11.5 min, 2%-99% B; 11.5−15 min, 99% B; 15−15.5 min, 99−0% B; 15.5−20 min, 0% B. For analysis of permethylated ribonucleosides, the following binary gradient was used: 0−7 min, 20%-40% B; 7−10 min, 40−70% B; 10−11 min, 70%-99% B; 11−15 min, 99% B; 15−16 min, 99−20% B; 16−20 min, 20% B. Samples were introduced into the mass spectrometer using electrospray ionization (ESI) source operated in the positive ion mode at 3500 V. Nitrogen sheath gas, auxiliary gas, and sweep gas flow rates were set at 30, 5, and 2 psi, respectively. The ion transfer tube temperature and vaporizer temperature were set at 350 °C and 175 °C, respectively. The precursor ions were fragmented using collision-induced dissociation (CID) with optimized energy. Data acquired from the UHPLC/QqQ-MS was collected using Thermo Scientific Xcalibur software (v4.1), and data analysis was performed using Thermo Scientific FreeStyle software (v2.1).

### nanoLC-MS/MS analysis

The samples were characterized using an EASY-nLC™ 1200 system coupled with a Q Exactive mass spectrometer (ThermoFisher Scientific). 3 μL of the sample was injected, and the analytes were separated on self-packed C18 column (3 μm, 0.150 mm × 250 mm) at a flow rate of 700 nL/min. Water containing 0.1% formic acid and 80% acetonitrile containing 0.1% formic acid were used as solvents A and B, respectively. MS spectra were collected with the mass range 200–600 m/z in positive ionization mode. The filtered precursor ions in each MS spectrum were fragmentated via high collisional dissociation (HCD) at 30% normalized collision energy (NCE) with nitrogen gas.

## Results and discussions

### Optimization of Permethylation Reaction

During the process of permethylation, all hydrogens on hydroxyl, amine, and carboxyl groups in ribonucleoside molecules are replaced with methyl groups. Conventionally, permethylation with DMSO and NaOH in solution is used, but the reaction efficiency hindered the application of this derivatization approach. Kang *et al*. demonstrated a solid-phase permethylation technique by packing sodium hydroxide beads in microspin columns.^33^ High derivatization efficiency was achieved in a short time using this technique and it has been extensively applied to improve the characterization of glycans and glycoproteins. Therefore, we employed this solid-phase-based technique to maximize the permethylation efficiency for the ribonucleoside samples (**Figure 1a**). The amount of water is critical for the permethylation reaction. To determine the optimal condition, the reaction was carried out on a mixture of four canonical ribonucleoside standards (2.5 μg/mL each) using varying volumes of water. Reactions performed in presence of 1 μL of water produced more permethylated ribonucleosides compared to reactions with 0.5 μL of water and 2 μL of water (reaction 1-3 in **Figure S1a**). We speculate that water improved the solubility of ribonucleosides in DMSO, while the extra amount of water led to undesired side reactions resulting in reduced efficiency of the permethylation. In addition, Mechref and co-workers demonstrated that reaction efficiency could be improved by adding extra iodomethane in the middle of the reaction.^34^ As the reaction 4 shows in **Figure S1a**, the yields of permethylated ribonucleosides increased dramatically and reached a maximum after the second aliquot of iodomethane-*d*_3_ was added to the reaction mixture.

**Figure 1.**
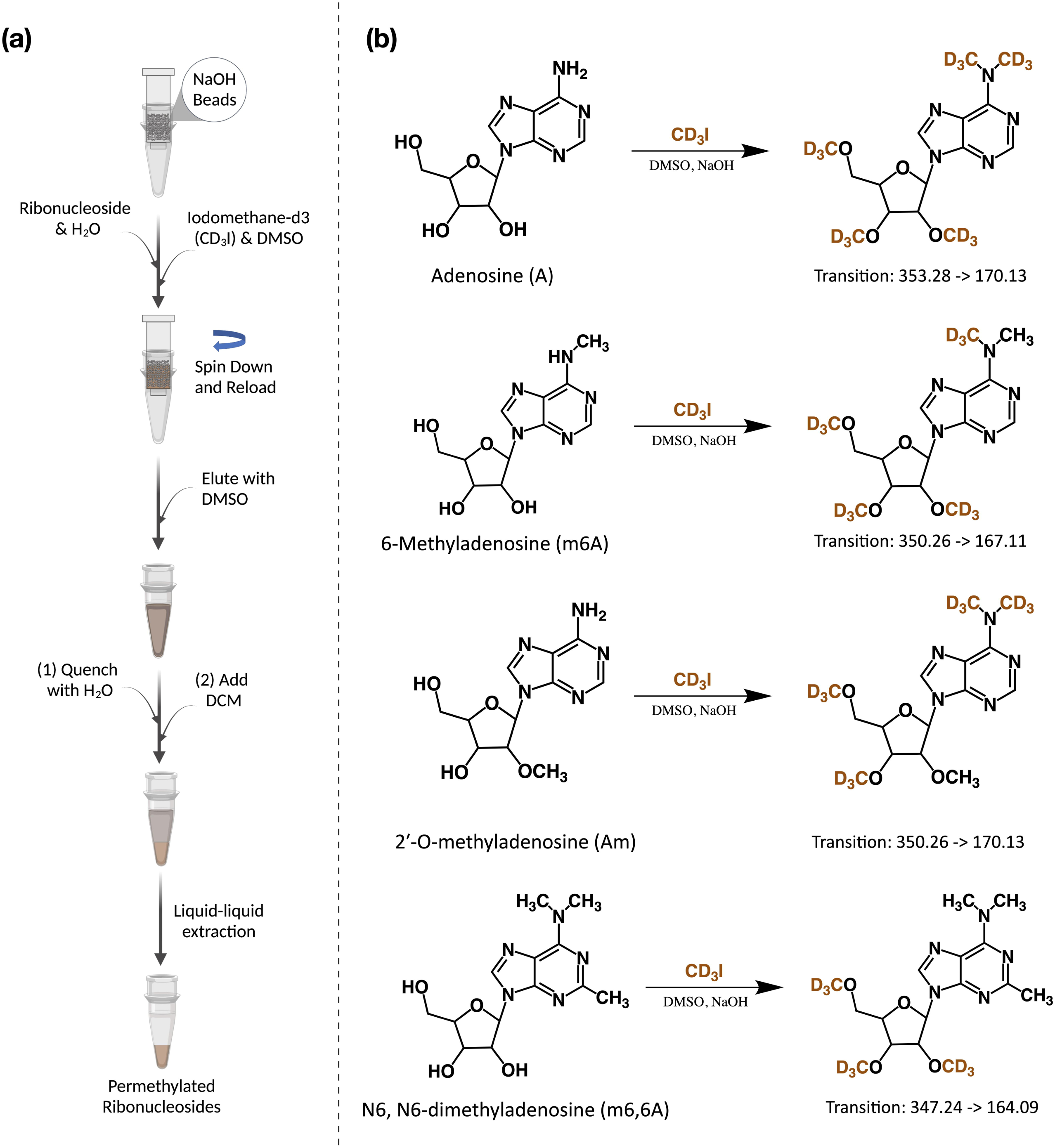
Permethylation was used to derivatize the ribonucleosides. **(a)** The workflow of the solid-phase permethylation method. **(b)** The example of permethylation of unmodified and methylated adenosine. Adenosine, 6-methyladenosine, and 2’-*O*-methyladenosine were labeled by the -CD_3_ group differently, and their unique precursor and product ions were monitored using UHPLC-QqQ-MS under the dMRM mode.

We also monitored the unreacted ribonucleosides in the mixture that were extracted from the aqueous phase during the liquid-liquid extraction (**Figure S1b**). The results showed that there were limited unreacted ribonucleosides after the reaction in all four conditions, and the unreacted species were even less abundant after adding the second aliquot of iodomethane-d_3_ (less than 0.005% for adenosine). Furthermore, the permethylation reaction may not fully replace all the active hydrogens, generating partially methylated products, which would hamper accurate quantification. For example, adenosine was noted to have an incomplete derivatization product using the conventional in-solution method.^35^ To ensure that the permethylation goes to completion, we used LC-MS/MS to characterize the adenosine products after solid-phase derivatization. As shown in Figure S2, the signal of fully permethylated adenosine was exceedingly abundant, while the partially methylated adenosine abundance was at the noise level. This result demonstrated the high efficiency of the optimized solid-phase permethylation method (>99.9%). To this end, 1 μL of water and the post-reloading supplemental aliquot of 20 μL of iodomethane-d_3_ was utilized for the complete derivatization of the ribonucleosides in our experiments.

### Construction of the dMRM Transitions

Commercial ribonucleoside standards were employed to create the basic transitions. Standards that were used to construct the transitions include A (adenosine), Am (2’-O-methyladenosine), m6A (N6-methyladenosine), t6A (N6-threonylcarbamoyladenosine), io6A (N6-(cis-hydroxyisopentenyl)adenosine), i6A (N6-isopentenyladenosine), I (inosine), C (cytidine), ac4C (N4-acetylcytidine), s2C (2-thiocytidine), m5C (5-methylcytidine), f5C (5-formylcytidine), G (guanosine), m7G (7-methylguanosine), U (uridine), m5U (5-methyluridine), s2U (2-thiouridine), D (dihydrouridine), m5D (5-methyldihydrouridine), and Ψ (pseudouridine). For analysis, permethylated ribonucleoside standards were prepared, and dMRM transitions of the permethylated standards were obtained by scanning their respective fragment ions using the product ion mode of the QqQ.

To distinguish the endogenous methylated molecules from the methylated molecules after derivatization process, deuterium-labeled iodomethane was used for the reaction. As shown in **Figure 1b**, five *d*_3_-methyl (−CD_3_) groups replaced hydrogen atoms from A, including three from the hydroxyl group on ribose ring and two from the amine group on nucleobase, while m6A and Am were labeled with four molecules of -CD_3_. Notably and as expected, -CD_3_ groups labeled hydroxyl and amine groups on m6A and Am differently. Two molecules of -CD_3_ on the ribose and two on the nucleobase replaced hydrogens for Am, while three hydrogens on ribose and one hydrogen on nucleobase were replaced by -CD_3_ for m6A. Only the hydrogens on ribose were replaced by -CD_3_ for N6, N6-dimethyladenosine (m6,6A). Similar to the underivatized form, the primary fragmentation of these permethylated ribonucleosides was produced by ribose ring loss, hence, three different transitions were able to be constructed, *m/z* 353.28 → 170.13 for A, *m/z* 350.26 → 167.11 for m6A, *m/z* 350.26 → 170.13 for Am, and *m/z* 347.24 → 164.09 for m6,6A. Notably, the permethylation product of m6A and m1A had the same mass with *m/z* 350.26. To distinguish these two isomers, a further (pseudo)-MS^3^ is required and the methyl position can be differentiated with different abundance distribution at m/z 119.03 and m/z 120.04 (**Figure S3**). Importantly, permethylation yielded the unmodified and methylated ribonucleosides with similar structures which have the same retention time, allowing the transition list to be built more readily. For example, after the transition and retention time of inosine were monitored based on its standard, the other methylated species, including Im, m1I, and m1Im, could also be spontaneously created without those standards.

In addition, the permethylation also improved the ability to distinguish isomers. For example, the isomer of canonical uridine, Ψ, is a crucial RNA modification and is involved in regulation of gene expression.^36^ Different techniques have been developed for the analysis of Ψ, but the same precursor mass and poor separation of the two isomers using RP-LC have hindered their analysis. Notably, these two isomers generated two different precursor and product ions after derivatization, which suggests that U and Ψ can be unambiguously characterized using our method. As mentioned above, most of the primary fragmentation of ribonucleosides came from ribose ring loss; however, Ψ was an exception due to the carbon-carbon linkage between the ribose and the nucleobase. In order to find the fragments of Ψ, we analyzed permethylated U and Ψ standards using a high-resolution Orbitrap MS. As shown in **Figure 2a**, U lost the methylated ribose ring (−180.14) and yielded its unique reporter ion of *m/z* 130.07. Interestingly, *m/z* 149.11 was the most abundant fragment for U, while it was not selected as the product ion. This is because it was generated from the cross-ring fragmentation of permethylated ribose and could also be produced by all other permethylated ribonucleosides. Additionally, the fragment ions of *m/z* 206.15 and 225.11 were uniquely produced by Ψ due to the cross-ring fragmentation (**Figure 2b**). Therefore, the transitions *m/z* 313.22 → 130.07 and *m/z* 330.25 → 206.15 were created for U and Ψ, respectively. It is also challenging to distinguish the two methylated uridine isomers, m5U and m3U, using the conventional method. After permethylation, m5U had one more -CD_3_ molecule labeled compared to m3U, which yielded differences in precursor ion mass, product ion mass, and retention time.

**Figure 2.**
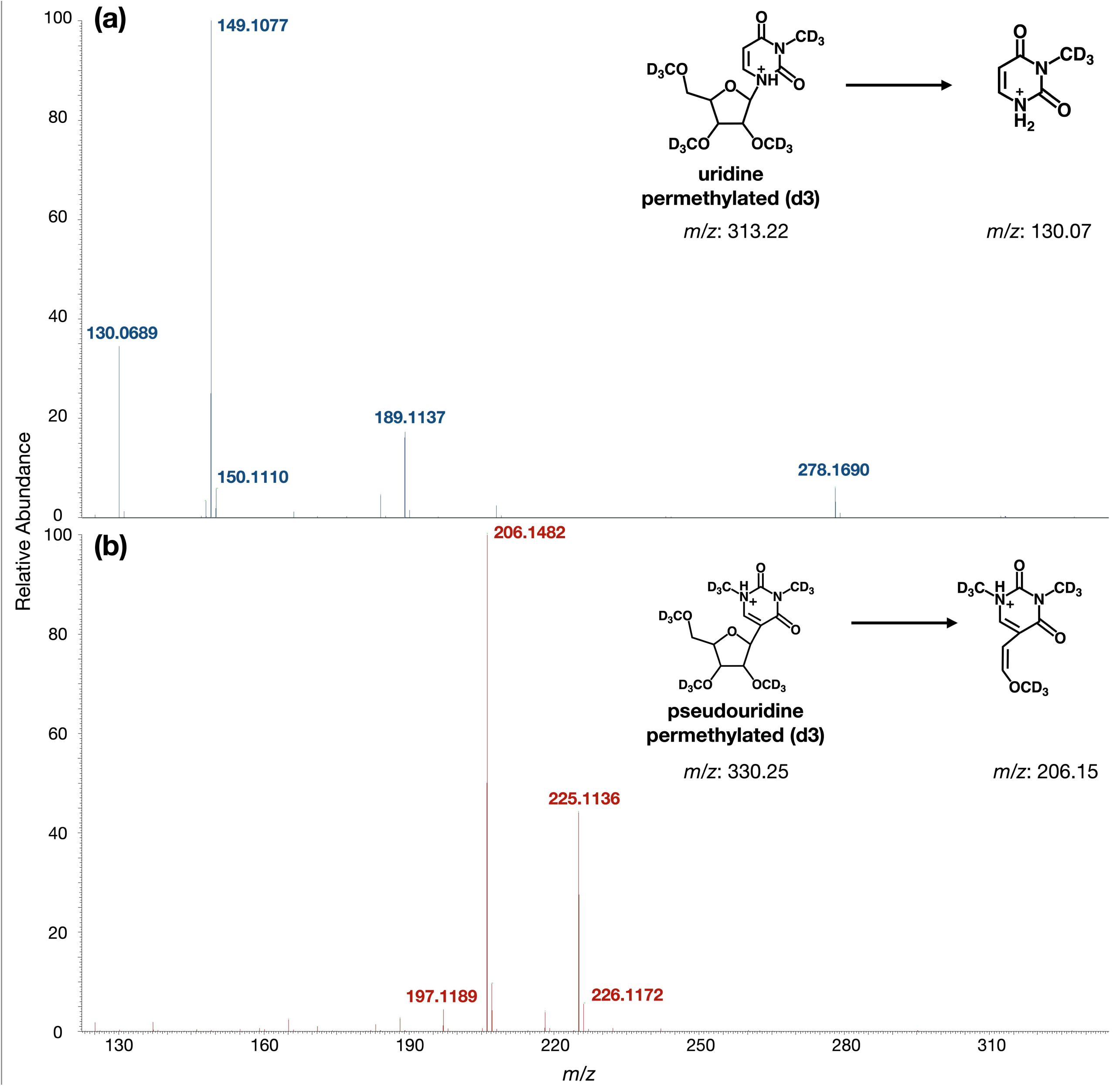
LC-MS/MS spectra of two permethylated isomers: uridine (U) and pseudouridine (Ψ). MS/MS spectra of the methylated(*d*_3_)-labeled protonated precursor ion (a) *m/z* 313.22 for U and (b) *m/z* 330.25 for pseudouridine Ψ. Fragmentation by HCD of U resulted in signals m/z 149.11 and m/z 130.07 as the two most abundant fragment ions, while the fragmentation of Ψ produced signals m/z 206.15 and m/z 22.5.11 as the two most abundant fragment ions.

We further monitored three adenosine-derived ribonucleoside standards i6A, io6A, and t6A, which have more complicated nucleoside base structures than methylated adenosine. Similar to the unmodified adenosine, the -CD_3_ groups labeled hydroxyl groups in the ribose and amines in the nucleobase of i6A (**Figure S4a**), while an extra -CD_3_ group reacted with io6A due to its hydroxyisopentenyl group (**Figure S4b**). Although permethylated io6A also generated the i6A ion in high intensity due to fragmentation of the -OCD_3_, both standards still generated high abundance fragments from ribose loss. Threonylcarbamoyl-modified adenosine, t6A, contains an acidic structure. The carboxyl group reacted with iodomethane-d_3_ and generated a product with *m/z* 532.38 as the precursor mass, and the expected fragment at *m/z* 349.23 was monitored in positive ion mode (**Figure S4c**). One exception is positively charged m7G, where a hydroxyl group was added from the hydroxide (**Figure S5a**). As a result, one extra methoxy group was yielded, and m7G was monitored using *m/z* 435.37 as the precursor ion and *m/z* 252.22 as the product ion (**Figure S5b**). In general, same as the glycans, -CD_3_ group can completely replace all the hydrogens on hydroxyl, amine, and carboxyl groups for ribonucleosides. This suggested that the products of derivatization are able to be easily monitored for structurally complex RNA modifications. Overall, more than 60 transitions were monitored simultaneously and shown in **Supplementary Table 1**, and the qualifying fragment ions of the permethylated ribonucleosides were also chosen to validate the method.

### Optimization of Collision Energies

For underivatized ribonucleosides, low collision energy (∼10 eV) was utilized for the identification and quantification due to the unstable nature of the ribonucleosides. Because the permethylation can enhance the stability of the precursor ions in the gas phase, the energy required for the fragmentation needed to be optimized. Therefore, we optimized the collision energies ranging from 10 to 35 eV for the permethylated ribonucleosides to obtain the best signal for the quantifying fragment ions. As shown in **Figure S6**, responses of most permethylated ribonucleosides first increased with rising collision energy, caused by more efficient fragmentation of the precursor ions. Signals decreased when the collision energy was set greater than 20 eV, which may be due to the larger percentage of over-fragmentation under the high-energy collision conditions. As a result, we found that 15-20 eV yielded the best signals for the majority of the permethylated ribonucleosides.

### Comparison with conventional ribonucleoside analysis

The ribonucleosides have similar structures, which contain hydrophilic ribose and a nucleobase. Therefore, it is a challenge to profile and quantify these compounds by traditional RP-LC-MS methods. To evaluate the advantages of the derivatized ribonucleosides compared to the underivatized form, equal amounts of standards were injected into the instrument and analyzed. As shown in **Figure 3a** and **b**, all four unpermethylated ribonucleoside standards were eluted within 4 mins under 1% of ACN. Notably, the chromatogram of these ribonucleosides had problems in peaks overlapping during the analysis. For example, the more hydrophilic ribonucleosides, cytidine and uridine had poor retention on C18 and were eluted in 0% ACN, which risks co-elution with hydrophilic contaminants. On the other hand, although purines including adenosine and guanosine retained on the reversed-phase, they bound weakly, so they also had poor chromatographic resolution. Ionization suppression therefore can consequently reduce sensitivity of hydrophilic ribonucleosides, so separation of the nucleosides is critical. Meanwhile, all the permethylated ribonucleoside peaks were evenly distributed across the chromatogram with a higher percentage of ACN (∼30% ACN) (**Figure 3c** and **d**), which indicated the chromatographic resolution of ribonucleosides on C18 was improved after labeling with the -CD_3_ groups. The theoretical plate number was calculated based on the retention time and the full peak width and shown in **Supplementary Table 2**. As a result, the number of theoretical plates for conventional ribonucleoside analysis was around 4000, while it was greatly improved for permethylated ribonucleosides even when a much higher percentage of ACN was applied. For example, the theoretical plate number was more than 26000 for permethylated guanosine, and nearly 16000 for permethylated adenosine. This demonstrated that C18 can continue to be the column resin of choice for nucleoside analysis without suffering poor resolution and analyte retention. Therefore, permethylation would allow users to avoid cumbersome HILIC chromatography for the analysis of RNA modifications.

**Figure 3.**
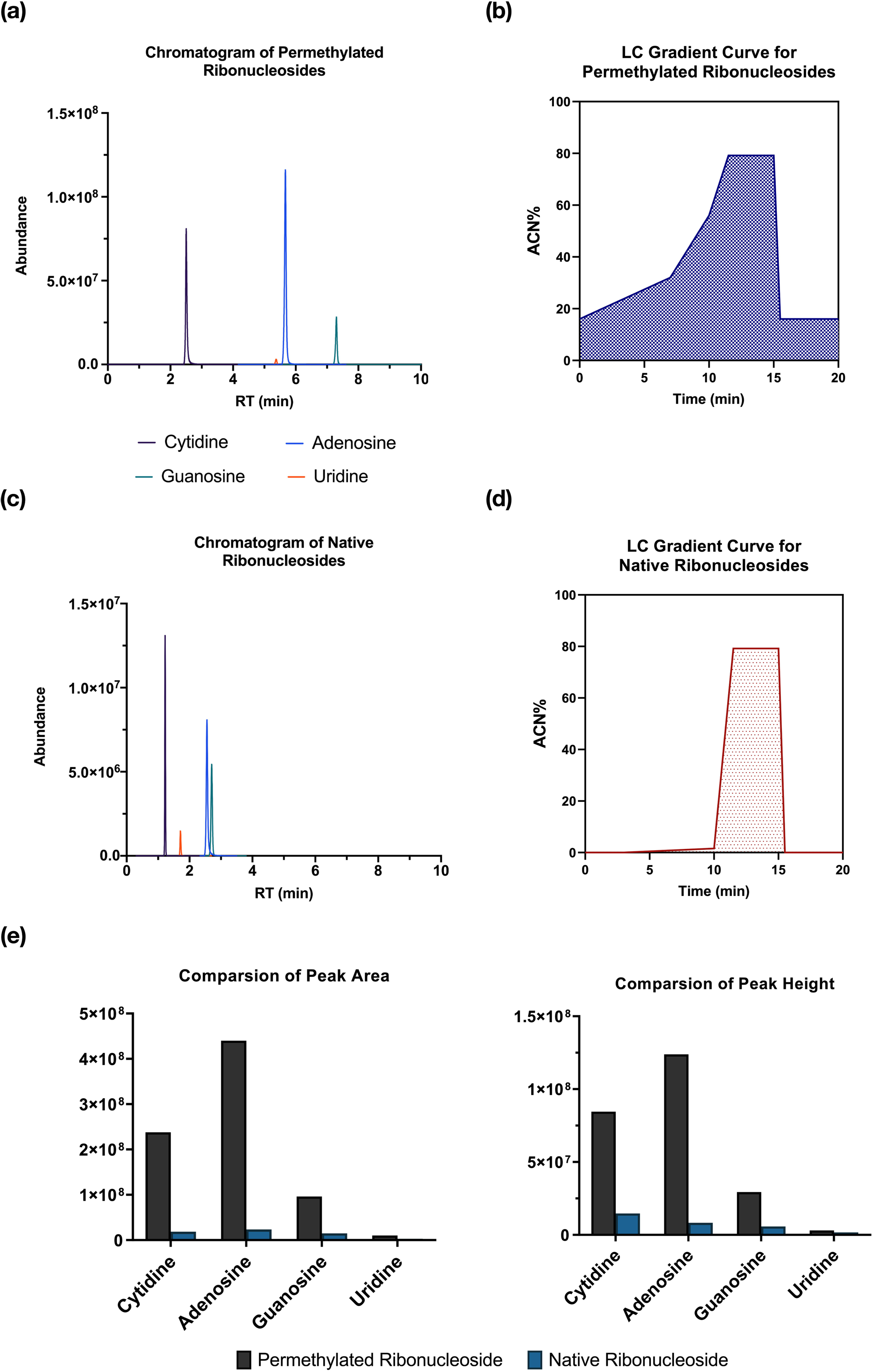
Comparison of the analysis between permethylated and native ribonucleoside. MRM chromatogram of. **(a)** permethylated(*d*_3_) ribonucleoside standards (c) native ribonucleoside standards. The LC gradient was applied for analyzing (b) permethylated(*d*_3_) canonical ribonucleosides and (d) the native forms. The ACN% (B%) w much higher for permethylated(*d*_3_) ribonucleosides analysis and the separation of analytes was improved. (e) The comparison of peak area and peak height between permethylated and native ribonucleosides.

Due to the hydrophilicity of the ribonucleosides, the compounds normally had low positive mode ionization efficiencies that resulted in difficulties in quantitative measurement.^37^ It has been demonstrated that permethylation can enhance the signals of hydrophilic analytes.^38^ Indeed, the signal of permethylated ribonucleosides was largely boosted compared to the underivatized ribonucleosides (**Figure 3 e** and **f**). For example, the peak area of the adenosine increased as much as 20-fold. This is likely due to the high hydrophilicity of underivatized ribonucleosides leading to lower proton affinity. At the same time, the incorporation of a large number of methyl groups with increased hydrophobicity can provide higher ionization efficiency in positive electrospray ionization. Uridine still had the lowest ionization efficiency among four canonical ribonucleosides. Nonetheless uridine signal after the permethylation increased 5-fold compared to the unpermethylated form, which is particularly valuable for analyzing uridine-based ribonucleosides that are normally difficult to detect. In general, the process of permethylation improved ribonucleoside quantitative analysis with great retaining performance, better separation, and higher sensitivity.

### Method validation

Quantitative analysis of canonical and modified ribonucleosides was performed to validate the derivatization method. The standard ribonucleoside mixtures were prepared by mixing the standards together and diluting solutions to the concentration range of 0.01–1000 ng/L. Working solutions with different dilution factors were derivatized according to the optimized solid-phase permethylation workflow. The calibration curve, linear regression coefficient (R^2^), linear range, limits of detection (LOD), limits of quantification (LOQ), and coefficients of variance (CV) for the permethylated ribonucleosides were calculated and shown in **Supplementary Table 3**. As a result, adenosine had the highest responses among all the standards due to the higher ionization efficiency of the molecule. All the correlation coefficients for the analytes were between 0.992-0.998, which indicated very good linearity of this method. Besides, both LOD and LOQ values were mostly at sub-femtomole levels, which was one order of magnitude lower compared to the analysis for underivatized ribonucleosides.^39^ Notably, LOD and LOQ for dihydrouridine and 5-methyldihydrouridine were higher than other ribonucleosides, which might be because the dihydrouridine is sensitive to the alkaline conditions wherein the pyrimidine ring opened and yielded by-products.^40^ This result suggested that this method is suitable for analyzing broad RNA modifications found in low abundance in biological samples. Furthermore, the linear calibration ranges for the permethylated ribonucleosides demonstrated good linearity within three orders of magnitude, while the technical replicate CVs were nearly 4% for most ribonucleosides. Collectively, this method showed an improved quantitative analysis compared to the ribonucleoside analysis without permethylation.^21^

### Analysis of ribonucleosides from mESCs

To demonstrate the applicability of the method, we quantified the levels of C, Cm, hm5C, and m5C in RNAs extracted from mESCs carrying mutations in genes encoding TET dioxygenases as well as wildtype (WT) controls. TET dioxygenases, which include TET1, TET2, and TET3, play important roles in epigenetic regulation because they catalyze the oxidation of 5-methylcytosine in DNA to form oxidized products.^41^ Recently, He et al. demonstrated that TET2 is also necessary for the conversion of m5C to hm5C on tRNAs, which regulates the formation of tRNA fragments.^32^ In this previous study, hm5C was measured using native ribonucleoside MS analysis and was shown to be substantially decreased in TET mutants. However, our previous analyses were not sufficiently accurate to detect the expected corresponding increase in m5C.

We repeated these measurements using our improved permethylation-based method (**Figure 4a** and **b**). Consistent with our published results, we detected a significant decrease in hm5C in RNA obtained from TET2-deficient mESCs, which was even more pronounced in RNA from triple *Tet1/2/3* knockout (KO) cells (**Figure 4c**).^32^ Notably, our improved method detected 10-fold higher levels of hm5C in these samples compared to our previous results based on native ribonucleoside analyses, likely due to improved sensitivity and CVs, as discussed above. We were also able to detect the expected increase in m5C in both *Tet2* KO and *Tet1/2/3* triple KO as compared to WT cells (**Figure 4d**). As a control, levels of Cm, which is not known to be targeted by TET enzymes, were not affected by the mutations (**Figure 4e**).

**Figure 4.**
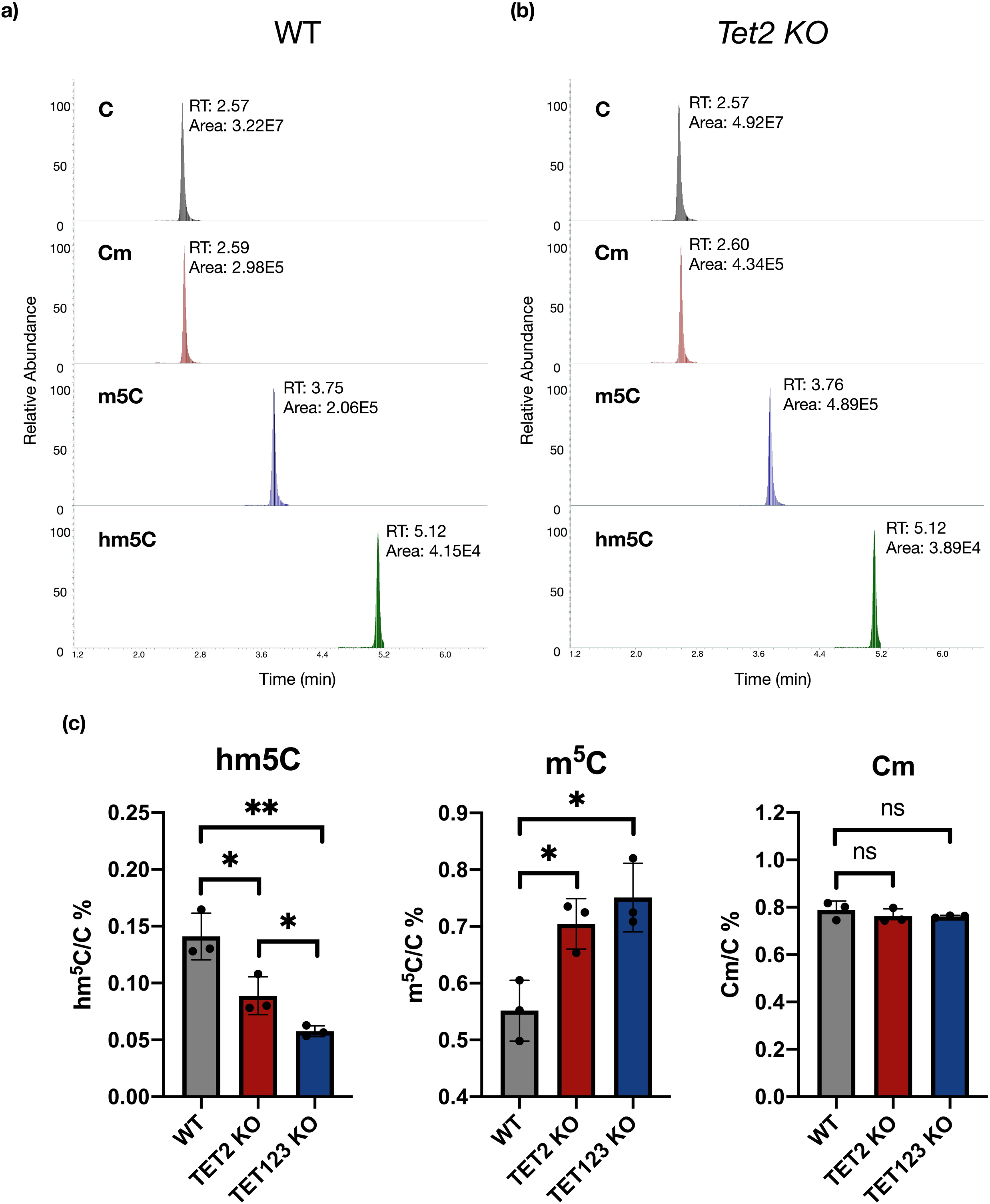
The application of the method for analyzing modified cytosine in WT and TET KO mESCs. The example chromatography of C, Cm, m5C, and hm5C in purified total RNAs under the condition of (a) WT and (b) TET2 KO. (c) The relative abundance of hm5C, m5C, and Cm. (p values were determined using two-tailed Student’s t test for unpaired samples. Error bars represent mean ± s.d., n = 3, ** p < 0.01, ***** p < 0.001, n.s. means p > 0.05.)

These observations were in agreement with our previous findings and provided more comprehensive information, including the level of m5C and Cm, simultaneously. Overall, these results exemplify the advantage of improved detection and quantification when performing the analysis of ribonucleoside modifications using the permethylation workflow.

## Conclusions

The analysis of the RNA modifications suffers from the hydrophilicity of the molecules, which can result in low sensitivity and inaccurate quantification. In this study, we derivatized ribonucleosides to hydrophobic molecules by using solid-phase permethylation. A dynamic MRM method was developed to provide better quantitative characterization for RNA modifications. The high reproducibility of the quantitative results suggests that this method essentially improves upon the conventional analysis and provides a powerful tool to monitor the ribonucleoside modifications on RNAs. Overall, this method is capable of profiling more than 60 modified ribonucleosides in RNAs with high throughput and the transition list can be readily extended as more RNA modifications are discovered. Similar to applications in glycan research, different isotopically labeled iodomethane can be further applied as multiplexing reagents to enhance the quantification of multiple samples simultaneously.^42^ This method can also be potentially employed for detecting canonical and modified deoxyribonucleosides in DNA samples, which provides a tool for facilitating both epigenetic and epitranscriptomic analysis. Additionally, the quantitative results of ribonucleosides extracted from WT and TET KO mESCs demonstrate that this approach can detect differences in low-abundant, but important, RNA modifications in biological samples. As ribonucleoside modifications are associated with several diseases, this method provides a new reliable platform for biomarker research and discovery.

## Author contributions

Y.X. and K.A.J. designed and performed the experiments, analyzed data, produced all figures, and drafted the manuscript. A.S. and R.B. prepared TET KO mESCs, extracted RNAs for the analysis, analyzed data, and edited the manuscript. R.B. supervised A.S. B.A.G. planned the overall experimental project and co-wrote the manuscript.

## Acknowledgements

The authors thank Dr. Juan J. Castillo and Siyu Chen (University of California, Davis) for their suggestions on permethylation reaction. Research reported in this publication was supported by grant to BAG from the NIH AI118891, CA196539 and AG031862.

