## Supplemental Figures and Tables for "Permethylation of ribonucleosides provides enhanced mass spectrometry quantification of post-transcriptional modifications"

**Supplementary Figure 1. Optimization of permethylation reaction.**

**Supplementary Figure 2. LC-MS chromatography of permethylated and partially methylated adenosine.**

**Supplementary Figure 3.** Permethylation reaction of m7G (7-methylguanosine).

**Supplementary Figure 4.** Tandem MS/MS spectra of adenosine modifications.

**Supplementary Figure 5.** Optimization of collision energy.

**Supplementary Figure 6.** Differentiation of m6A and m1A using MS3.

**Supplementary Table 1.** The dynamic MRM transitions for monitoring ribonucleosides.

**Supplementary Table 2.** Summary of the number of theoretical plates for underivatized and permethylated ribonucleoside analyses.

**Supplementary Table 3.** Summary of linearities of permethylated ribonucleoside standards.

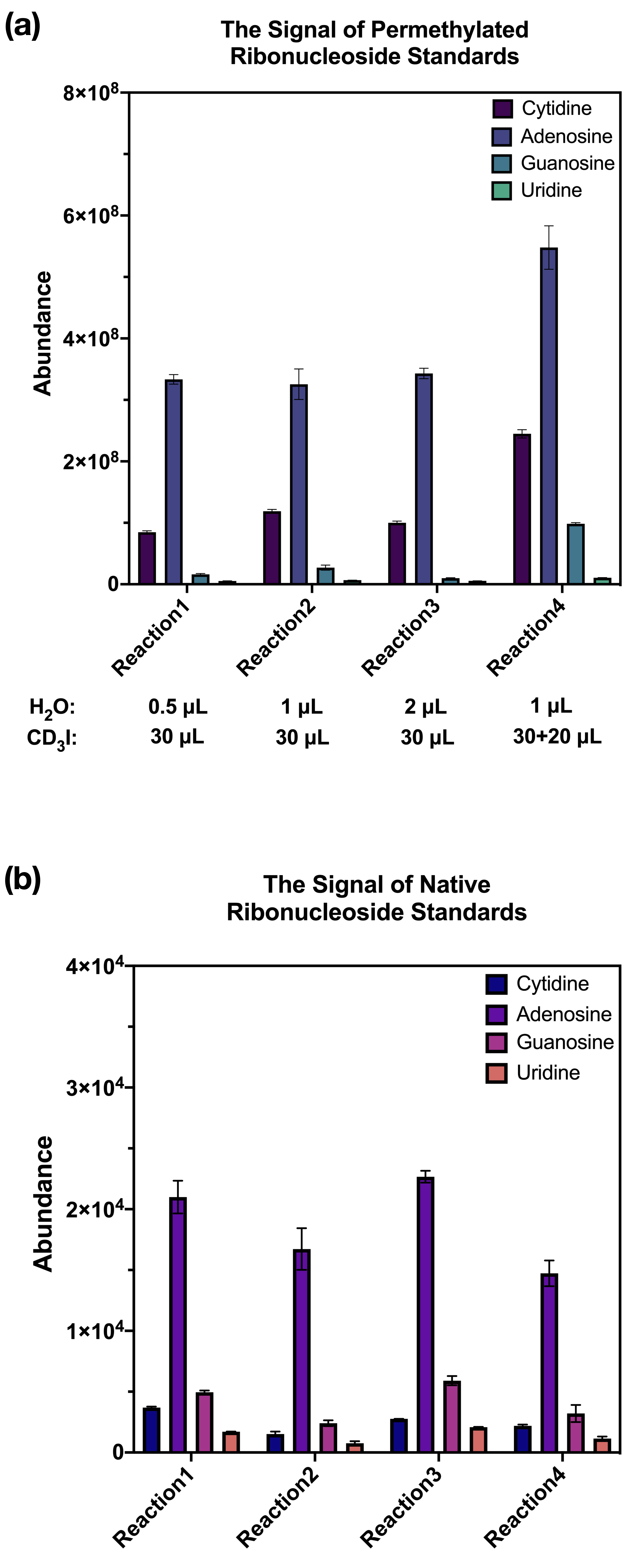

**Figure S1. Optimization of permethylation reaction.** The reaction was monitored by the signal of (**a**) permethylated ribonucleosides, and (**b**) unreacted ribonucleosides.

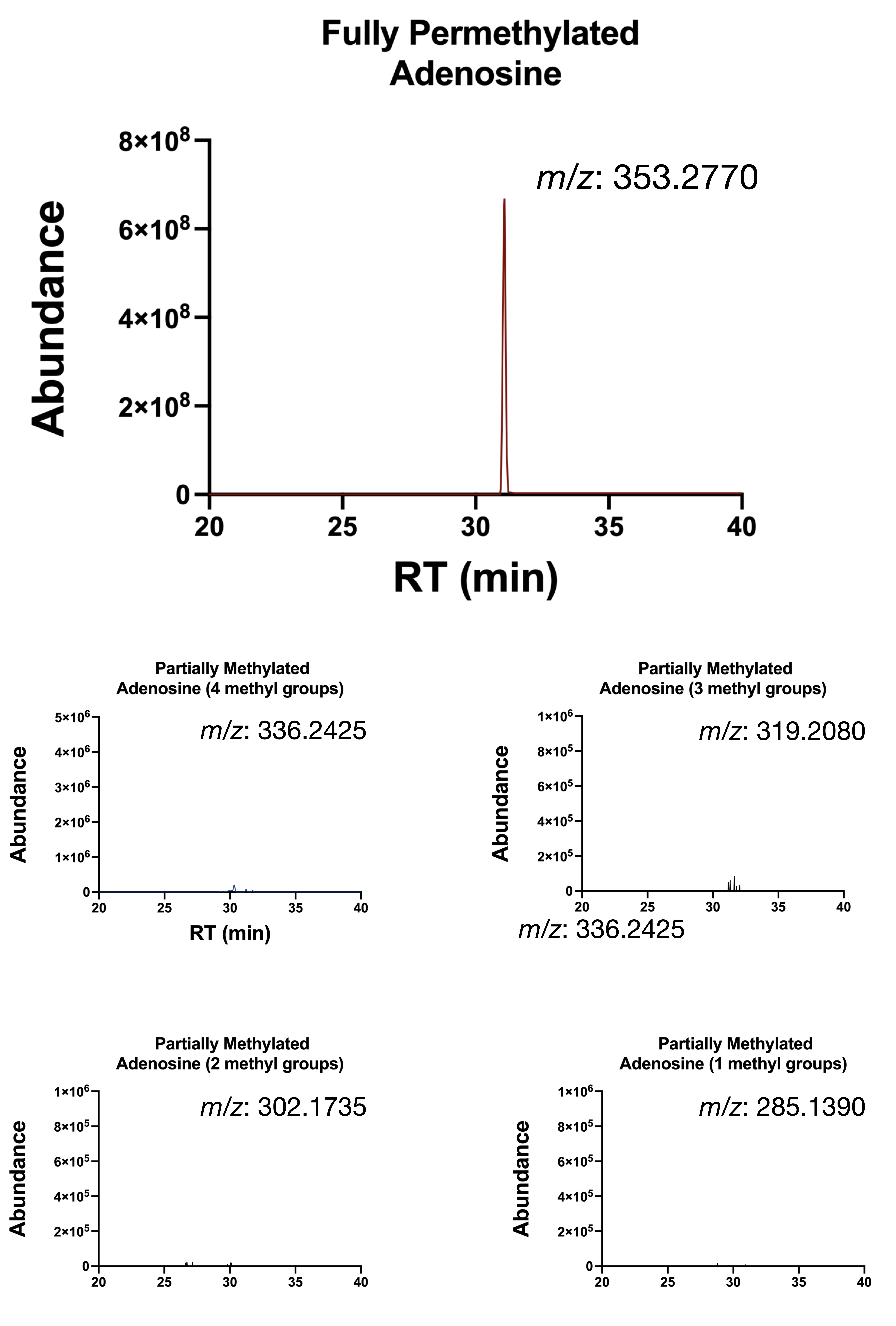

**Figure S2.** **LC-MS chromatography of permethylated and partially methylated adenosines.** Fully permethylated adenosine was monitored at *m/z* 353.2770, and four different partially methylated adenosines (with different methylation degrees) were monitored at *m/z* 336.2425, 319.2080, 302.1735, 285.1390, respectively.

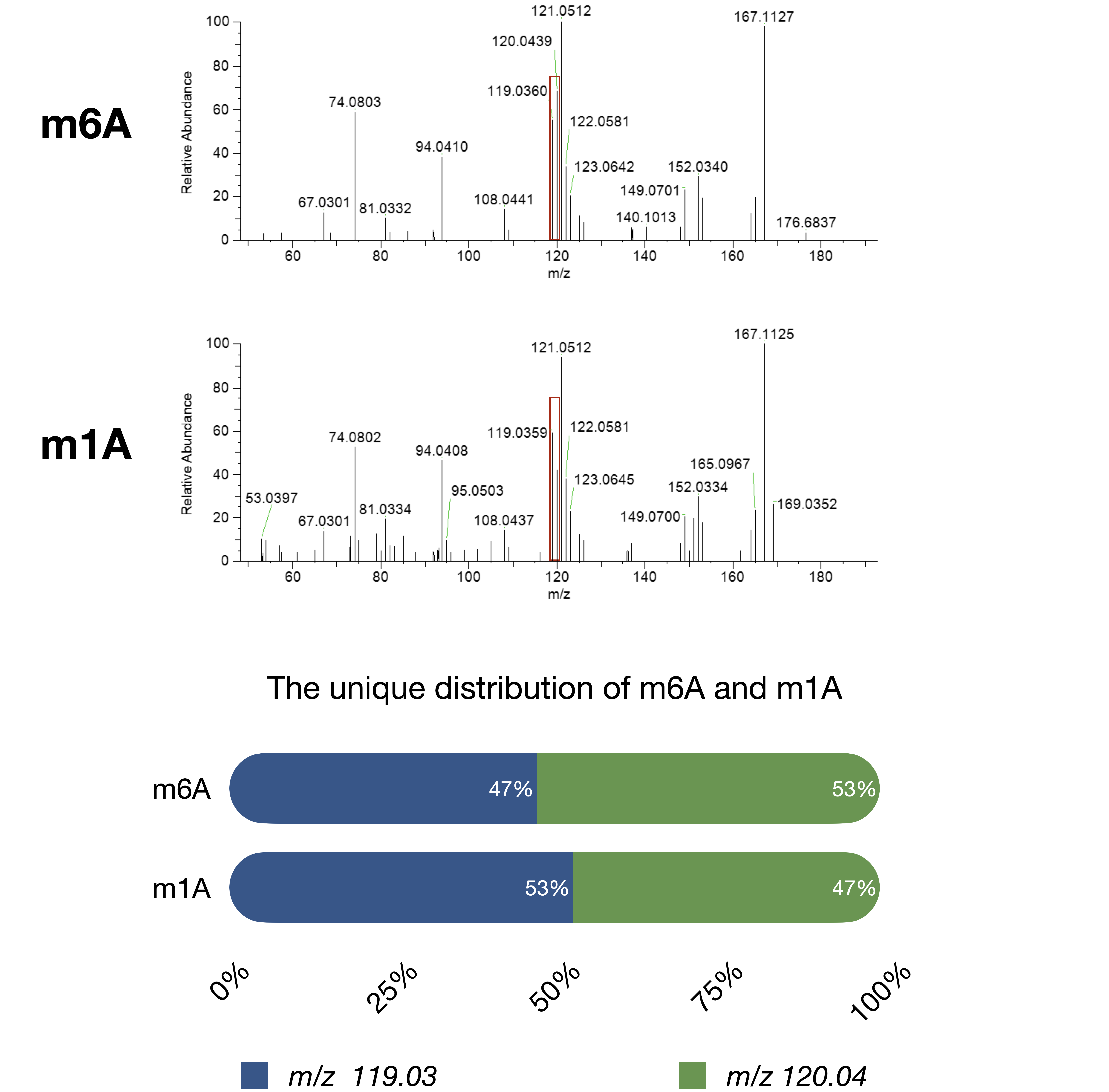

**Figure S3. Differentiation of m6A and m1A using MS3.** The fragmentation from nucleotide base, *m/z* 167.11, was selected for further fragmentation. The unique pattern at m/z 119.03 and m/z 120.04 could be used to distinguish m6A and m1A.

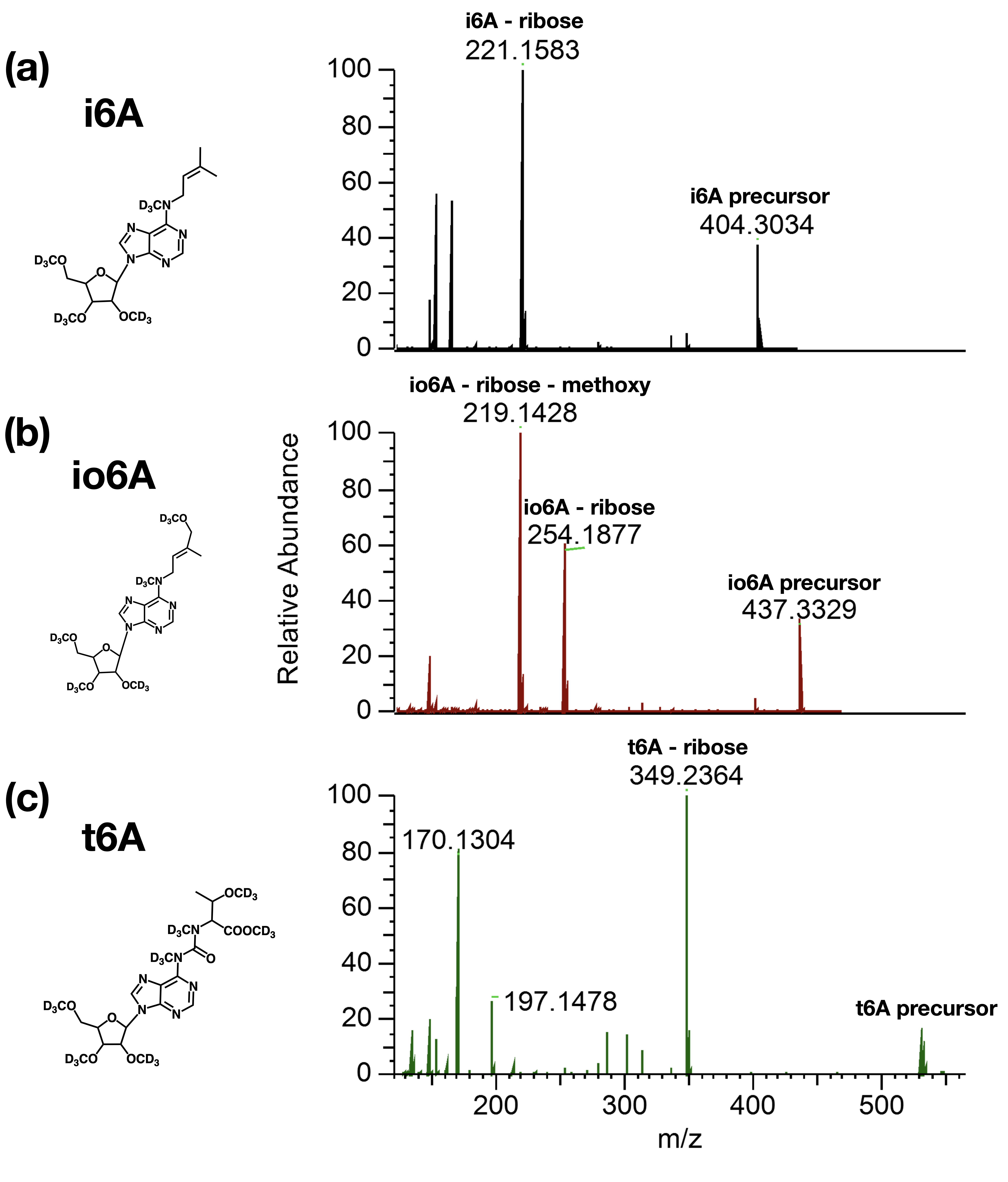

**Figure S4.** **Tandem** **MS/MS spectra of adenosine modifications.** The methylated(d3)-labeled (**a**) i6A, (**b**) io6A, and (**c**) t6A yielded their signature fragmentation patterns.

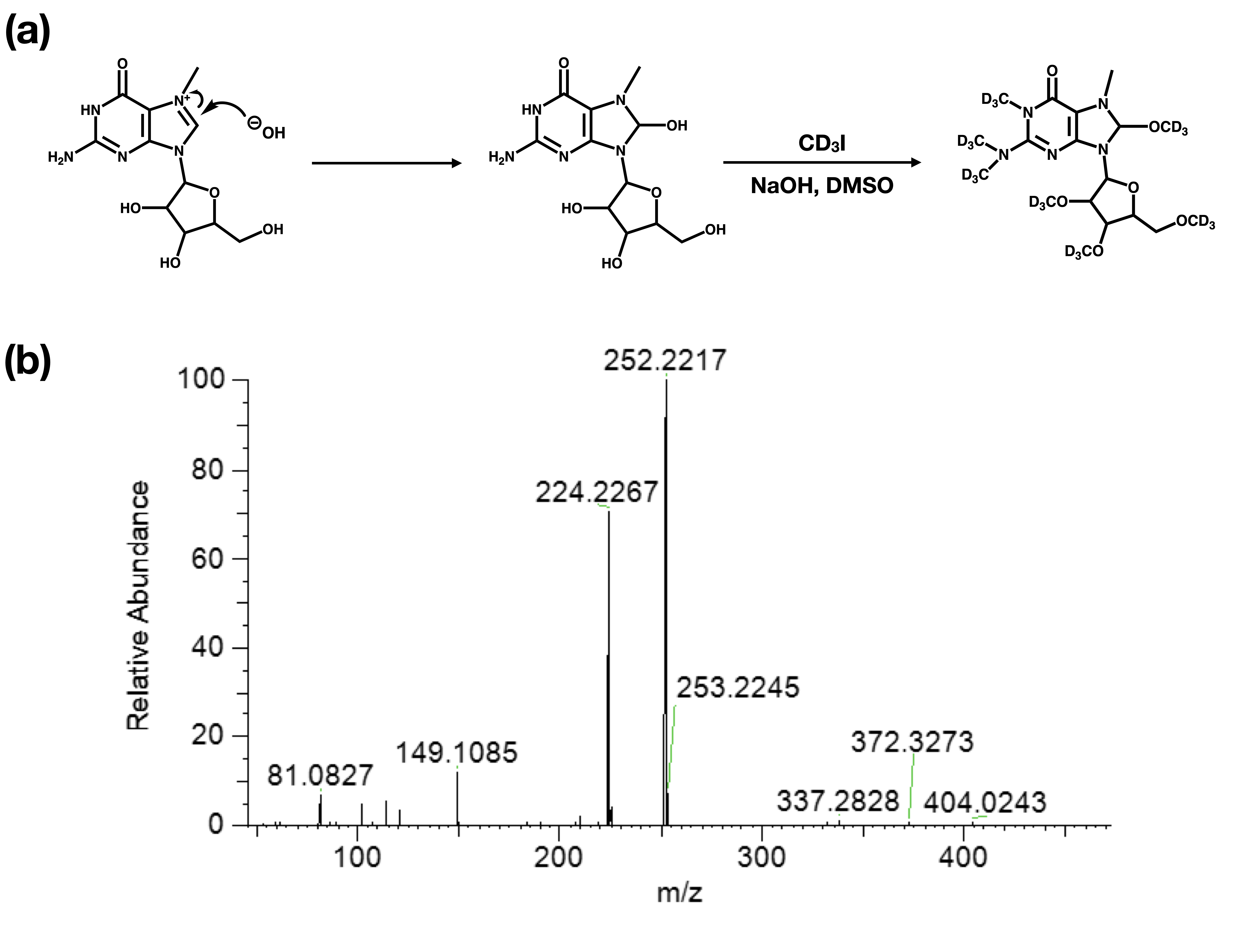

**Figure S5. Permethylation reaction of m7G (7-methylguanosine). (a)** Due to m7G being positively charged, an extra hydroxyl group was added. (**b**) The fragment at *m/z* 252.2217 was detected by monitoring the precursor ion of permethylated m7G at m/z 435.3671.

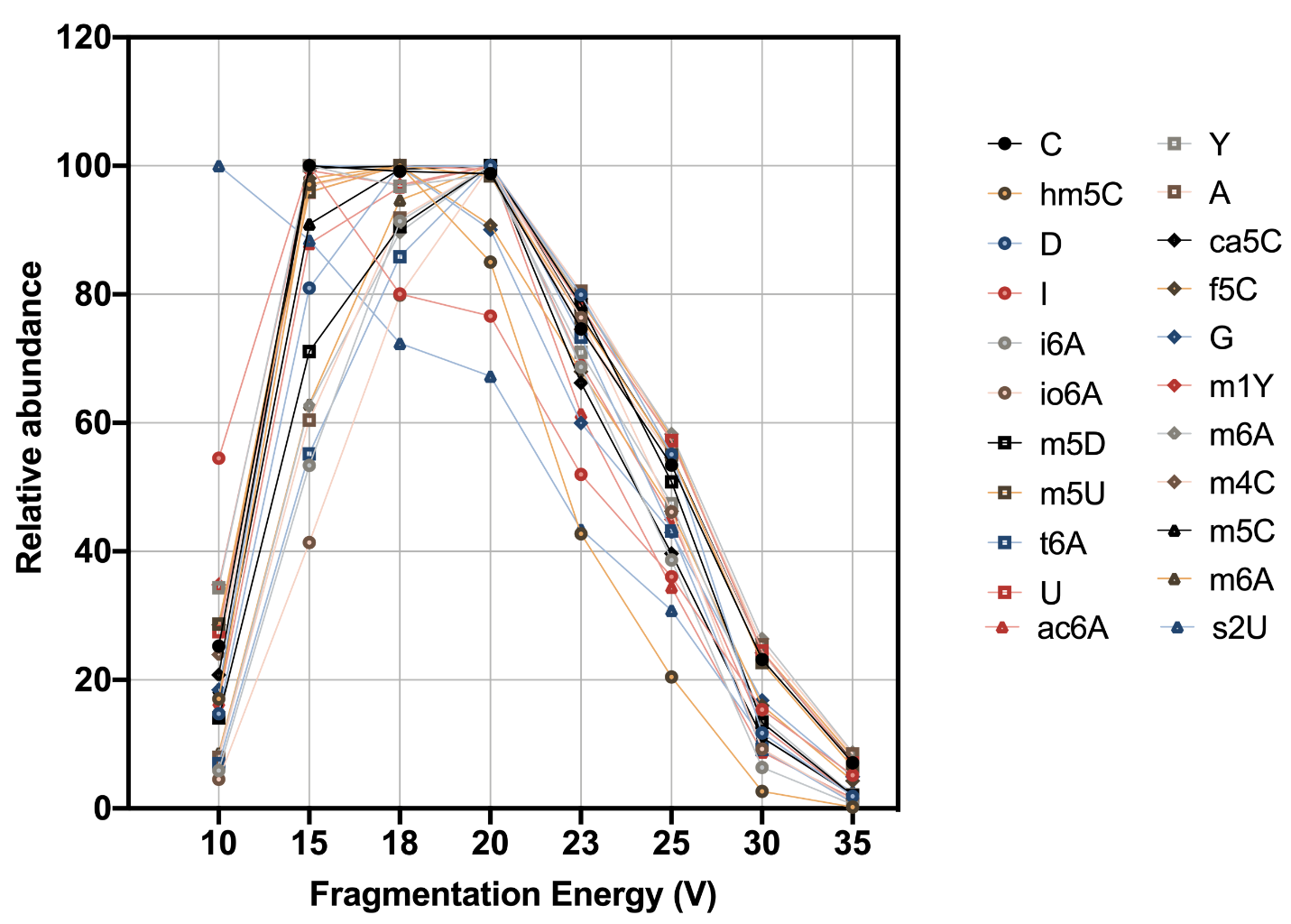

**Figure S6. Optimization of collision energy.** Permethylated ribonucleoside standards were fragmented under collision energies ranging from 10 to 35 eV to obtain the optimal signal for the analysis.

| **Compound Full Name** | **Common Name** | **Start Time**  **(min)** | **End Time**  **(min)** | **Precursor Ion**  **(m/z)** | **Product Ion**  **(m/z)** | **Collision Energy**  **(V)** |
| --- | --- | --- | --- | --- | --- | --- |
| 5-formylcytidine | f5C | 1.5 | 2.3 | 357.26 | 174.11 | 18 |
| 5-formyl-2'-O-methylcytidine | f5Cm | 1.5 | 2.3 | 354.24 | 174.11 | 18 |
| cytidine | C | 2.2 | 2.8 | 329.27 | 146.12 | 15 |
| 2'-O-methylcytidine | Cm | 2.2 | 2.8 | 326.25 | 146.12 | 15 |
| N4,N4-dimethylcytidine | m4,4C | 2.2 | 2.8 | 323.23 | 140.08 | 15 |
| N4,N4,2'-O-trimethylcytidine | m4,4Cm | 2.2 | 2.8 | 320.21 | 140.08 | 15 |
| N4-methylcytidine* | m4C | 2.2 | 2.8 | 326.25 | 143.1 | 15 |
| 3-methylcytidine* | m3C | 2.2 | 2.8 | 326.25 | 143.1 | 15 |
| N4,2'-O-dimethylcytidine* | m4Cm | 2.2 | 2.8 | 323.23 | 143.1 | 15 |
| 3-methyl-2'-O-methylcytidine* | m3Cm | 2.2 | 2.8 | 323.23 | 143.1 | 15 |
| 5-carboxylcytidine | ca5C | 2.54 | 3.14 | 390.29 | 207.14 | 18 |
| 5-carboxyl-2'-O-methylcytidine | ca5Cm | 2.54 | 3.14 | 387.28 | 207.14 | 18 |
| inosine | I | 3.3 | 3.9 | 337.23 | 154.08 | 15 |
| 2'-O-methylinosine | Im | 3.3 | 3.9 | 334.21 | 154.08 | 15 |
| 1-methylinosine | m1I | 3.3 | 3.9 | 334.21 | 151.06 | 15 |
| 1,2'-O-dimethylinosine | m1Im | 3.3 | 3.9 | 331.19 | 151.06 | 15 |
| 5-methylcytidine | m5C | 3.35 | 3.95 | 343.25 | 160.14 | 15 |
| 5,2'-O-dimethylcytidine | m5Cm | 3.35 | 3.95 | 340.26 | 160.14 | 15 |
| N4-acetylcytidine | ac4C | 3.35 | 4.15 | 354.24 | 171.1 | 15 |
| N4-acetyl-2'-O-methylcytidine | ac4Cm | 3.35 | 4.15 | 351.22 | 171.1 | 15 |
| 2-thiocytidine | s2C | 3.4 | 4 | 345.24 | 162.1 | 15 |
| dihydrouridine | D | 3.6 | 4.4 | 315.23 | 191.13 | 20 |
| 1-methylpseudouridine* | m1Y | 4.3 | 4.9 | 327.23 | 203.13 | 18 |
| 3-methylpseudouridine* | m3Y | 4.3 | 4.9 | 327.23 | 203.13 | 18 |
| pseudouridine | Y | 4.3 | 4.9 | 330.25 | 206.15 | 18 |
| 2'-O-methylpseudouridine | Ym | 4.3 | 4.9 | 327.23 | 206.15 | 18 |
| 5-hydroxymethylcytidine | hm5C | 4.6 | 5.3 | 376.31 | 193.16 | 18 |
| 2′‐O‐Methyl-5-hydroxymethylcytidine | hm5Cm | 4.6 | 5.3 | 373.29 | 193.16 | 18 |
| N6-acetyladenosine | ac6A | 4.9 | 5.5 | 378.25 | 195.11 | 20 |
| 3-methyluridine | m3U | 5.2 | 5.8 | 310.2 | 127.05 | 18 |
| 3,2'-O-dimethyluridine | m3Um | 5.2 | 5.8 | 307.18 | 127.05 | 18 |
| 2-thiouridine | s2U | 5.2 | 5.8 | 329.19 | 146.05 | 10 |
| 2-thio-2'-O-methyluridine | s2Um | 5.2 | 5.8 | 326.17 | 146.05 | 10 |
| uridine | U | 5.2 | 5.8 | 313.22 | 130.07 | 18 |
| 2'-O-methyluridine | Um | 5.2 | 5.8 | 310.2 | 130.07 | 18 |
| adenosine | A | 5.4 | 6 | 353.28 | 170.13 | 20 |
| 2'-O-methyladenosine | Am | 5.4 | 6 | 350.26 | 170.13 | 20 |
| 2,8-dimethyladenosine | m2,8A | 5.4 | 6 | 381.31 | 198.16 | 20 |
| 2-methyladenosine* | m2A | 5.4 | 6 | 367.29 | 184.15 | 20 |
| 8-methyladenosine* | m8A | 5.4 | 6 | 367.29 | 184.15 | 20 |
| N6,N6-dimethyladenosine | m6,6A | 5.6 | 6 | 347.24 | 164.09 | 20 |
| N6,N6,2'-O-trimethyladenosine | m6,6Am | 5.6 | 6 | 344.22 | 164.09 | 20 |
| N6-methyladenosine* | m6A | 5.6 | 6 | 350.26 | 167.11 | 20 |
| N6,2'-O-dimethyladenosine* | m6Am | 5.6 | 6 | 347.24 | 167.11 | 20 |
| 1-methyladenosine* | m1A | 5.6 | 6 | 350.26 | 167.11 | 20 |
| 1,2'-O-dimethyladenosine* | m1Am | 5.6 | 6 | 347.24 | 167.11 | 20 |
| 5-hydroxyuridine | ho5U | 5.85 | 6.45 | 346.24 | 163.1 | 18 |
| 5-methyldihydrouridine | m5D | 5.85 | 6.65 | 329.25 | 205.15 | 20 |
| 5-hydroxycytidine | ho5C | 6 | 6.6 | 362.3 | 179.15 | 15 |
| guanosine | G | 7.05 | 7.65 | 386.31 | 203.16 | 18 |
| 2'-O-methylguanosine | Gm | 7.05 | 7.65 | 383.29 | 203.16 | 18 |
| 1-methylguanosine* | m1G | 7.05 | 7.65 | 383.29 | 200.14 | 18 |
| 2-methylguanosine* | m2G | 7.05 | 7.65 | 383.29 | 200.14 | 18 |
| 1,2'-O-dimethylguanosine* | m1Gm | 7.05 | 7.65 | 380.27 | 200.14 | 18 |
| 2,2'-O-dimethylguanosine* | m2Gm | 7.05 | 7.65 | 380.27 | 200.14 | 18 |
| N2,N2-dimethylguanosine | m2,2G | 7.05 | 7.65 | 380.27 | 197.12 | 18 |
| N2,N2,2'-O-trimethylguanosine | m2,2Gm | 7.05 | 7.65 | 377.27 | 197.12 | 18 |
| 5-carbamoylmethyluridine | ncm5U | 7.2 | 7.8 | 404.23 | 221.09 | 18 |
| 5-carbamoylmethyl-2'-O-methyluridine | ncm5Um | 7.2 | 7.8 | 401.29 | 221.16 | 18 |
| 5-methyluridine | m5U | 7.35 | 7.95 | 327.23 | 144.09 | 18 |
| 5,2'-O-dimethyluridine | m5Um | 7.35 | 7.95 | 324.21 | 144.09 | 18 |
| N2,N2,7-trimethylguanosine | m2,2,7G | 7.4 | 8 | 429.33 | 246.18 | 25 |
| N2,7-dimethylguanosine | m2,7G | 7.4 | 8 | 432.35 | 249.20 | 25 |
| N2,7,2'-O-trimethylguanosine | m2,7Gm | 7.4 | 8 | 429.33 | 249.20 | 25 |
| 7-methylguanosine | m7G | 7.4 | 8 | 435.37 | 252.22 | 25 |
| N6-hydroxymethyladenosine | hm6A | 7.45 | 8.05 | 383.29 | 200.14 | 20 |
| 5-methoxycarbonylmethyluridine | mcm5U | 9.45 | 10.15 | 405.29 | 159.1 | 18 |
| 5-methoxycarbonylmethyl-2'-O-methyluridine | mcm5Um | 9.5 | 10.1 | 402.28 | 159.1 | 18 |
| N6-(cis-hydroxyisopentenyl)adenosine | io6A | 11.05 | 11.55 | 437.33 | 254.19 | 20 |
| N6-methyl-N6-threonylcarbamoyladenosine | m6t6A | 11.15 | 11.65 | 529.37 | 346.23 | 20 |
| N6-threonylcarbamoyladenosine | t6A | 11.15 | 11.65 | 532.38 | 349.24 | 20 |
| N6-isopentenyladenosine | i6A | 12.6 | 13.1 | 404.31 | 221.16 | 20 |
| 8-hydroxyguanosine | ho8G | 12.75 | 13.35 | 419.34 | 236.19 | 18 |

* The corresponding compounds have the same transition, the methyl position can be differentiated with MS^n^ fragmentation.

**Table 1. The dynamic MRM transitions for monitoring ribonucleosides.**

| Ribonucleoside | Retention time (min) | Peak Width (min) | Plate number (N) |
| --- | --- | --- | --- |
| C | 1.22 | 0.08 | 3721 |
| U | 1.71 | 0.1 | 4679 |
| A | 2.56 | 0.15 | 4660 |
| G | 2.71 | 0.16 | 4590 |
| C (permethylated) | 2.51 | 0.1 | 10080 |
| U (permethylated) | 5.38 | 0.16 | 18090 |
| A (permethylated) | 5.67 | 0.18 | 15876 |
| G (permethylated) | 7.3 | 0.18 | 26316 |

**Table 2.** **Summary of the number of theoretical plates for underivatized and permethylated ribonucleoside analyses.**

| Ribonucleoside | [Mass+H] | Calibration  Curve | Linear Range  (μg/ml) | LOD  (fmol) | LOQ  (fmol) | Linear Regression Coefficients  (R^2^) | CV  (%) |
| --- | --- | --- | --- | --- | --- | --- | --- |
| A | 353.28 | Y = 434596*X + 995861 | 0.0001-0.2 | 0.028 | 0.094 | 0.998 | 5.70 |
| G | 386.31 | Y = 154199*X + 402855 | 0.0001-0.2 | 0.026 | 0.086 | 0.998 | 2.09 |
| C | 329.27 | Y = 413520*X - 89569 | 0.001-0.2 | 0.030 | 0.101 | 0.998 | 6.51 |
| m5C | 343.25 | Y = 1317655*X + 7545963 | 0.001-0.2 | 0.029 | 0.097 | 0.998 | 3.11 |
| U | 313.22 | Y = 6470*X + 17004 | 0.001-0.5 | 0.319 | 1.064 | 0.998 | 4.82 |
| m5U | 327.23 | Y = 24213*X + 346904 | 0.001-0.5 | 0.306 | 1.019 | 0.996 | 4.39 |
| D | 315.23 | Y = 55.33*X + 457.3 | 0.02-0.5 | 6.344 | 21.149 | 0.998 | 6.18 |
| m5D | 329.25 | Y = 1161*X + 22377 | 0.02-0.5 | 6.074 | 20.248 | 0.995 | 3.02 |
| I | 337.23 | Y = 32018*X + 96970 | 0.0001-0.2 | 0.030 | 0.099 | 0.992 | 3.56 |
| Y | 330.25 | Y = 19897*X + 120645 | 0.0001-0.5 | 0.303 | 1.009 | 0.998 | 4.29 |
| hm5C | 376.31 | Y = 2533470*X + 21278753 | 0.0001-0.2 | 0.027 | 0.089 | 0.994 | 3.16 |
| s2U | 329.19 | Y = 5733*X + 4050 | 0.001-0.2 | 0.304 | 0.113 | 0.996 | 4.05 |
| io6A | 437.33 | Y = 195785*X + 619394 | 0.00001-0.2 | 0.023 | 0.076 | 0.995 | 3.36 |
| t6A | 532.38 | Y = 3490*X + 40604 | 0.0001-1 | 0.019 | 0.063 | 0.995 | 7.37 |
| i6A | 404.31 | Y = 371114*X + 2088335 | 0.0001-0.2 | 0.025 | 0.082 | 0.992 | 1.92 |
| f5C | 357.26 | Y = 49678*X - 1638376 | 0.001-1 | 1.3995 | 4.665 | 0.994 | 4.56 |

**Table 3. Summary of linearities of permethylated** **ribonucleoside standards.**
